# Effects of gap junction misexpression on synapses between auditory sensory neurons and the giant fiber of *Drosophila melanogaster*

**DOI:** 10.1101/331074

**Authors:** Sami H. Jezzini, Amelia Merced, Jonathan M. Blagburn

**Author notes:** Corresponding author: Jonathan M. Blagburn.

## Abstract

The synapse between auditory Johnston’s Organ neurons (JONs) and the giant fiber (GF) of *Drosophila* is structurally mixed, being composed of cholinergic chemical synapses and Neurobiotin-(NB) permeable gap junctions, which consist of the innexin Shaking-B (ShakB). Misexpression of one ShakB isoform, ShakB(N+16), in a subset of JONs that do not normally form gap junctions, results in their *de novo* dye coupling to the GF. This is similar to the effect of misexpression of the transcription factor Engrailed (En) in these same neurons, which also causes the formation of additional chemical synapses. In order to test the hypothesis that ShakB misexpression would similarly affect the distribution of chemical synapses, fluorescently-labeled presynaptic active zone protein (Brp) was expressed in JONs and the changes in its distribution were assayed with confocal microscopy. Both ShakB(N+16) and En increased the dye-coupling of JONs with the GF, indicating the formation of ectopic gap junctions. Conversely, expression of the ‘incorrect’ isoform, ShakB(N) abolishes dye coupling. However, while En misexpression increased the chemical contacts with the GF and the amount of GF medial branching, ShakB misexpression did not. ShakB immunocytochemistry showed that misexpression of ShakB(N+16) increases gap junctional plaques in JON axons but ShakB(N) does not. We conclude that both subsets of JON form chemical synapses onto the GF dendrites but only one population forms gap junctions, comprised of ShakB(N+16). Misexpression of this isoform in all JONs does not result in the formation of new mixed synapses but in the insertion of gap junctions, presumably at the sites of existing chemical synaptic contacts with the GF.

## Introduction

There are two types of synapse in the nervous system, chemical and electrical, which are molecularly and physiologically disparate. Chemical synapses involve an enormously complex array of presynaptic transmitter release mechanisms, postsynaptic receptors, and signaling pathways, whereas electrical are comprised of apparently simple cylindrical clusters of gap junctional proteins. In the past it was thought that these simpler electrical synapses were more abundant in ‘lower’ vertebrates and invertebrates, and were of lesser importance in mammals. It is now clear, however, that this is not the case [1,2], and that these two synapse types are often found together, located in close proximity and interacting functionally, in the adult brain as well as during development [2].

These ‘mixed’ chemical and electrical synapses are a typical feature of escape circuits in vertebrates and invertebrates [2], where the gap junctions are formed from connexin proteins, or the functionally analogous innexins, respectively [3]. In *Drosophila melanogaster* 8 genes for innexins have been identified [4]. One of the most thoroughly studied innexins, Shaking-B (ShakB), is a critical component of the giant fiber (GF) portion of the escape circuitry [5–7] (Fig. 1). ShakB-containing electrical synapses are present, along with cholinergic chemical synapses [8], at contacts made by the GF with two of its output neurons, allowing the fast activation of the ‘jump’ tergotrochanteral muscle (TTM) followed shortly by the dorsal longitudinal flight muscle (DLM) [9] (Fig. 1).

**Fig 1.**
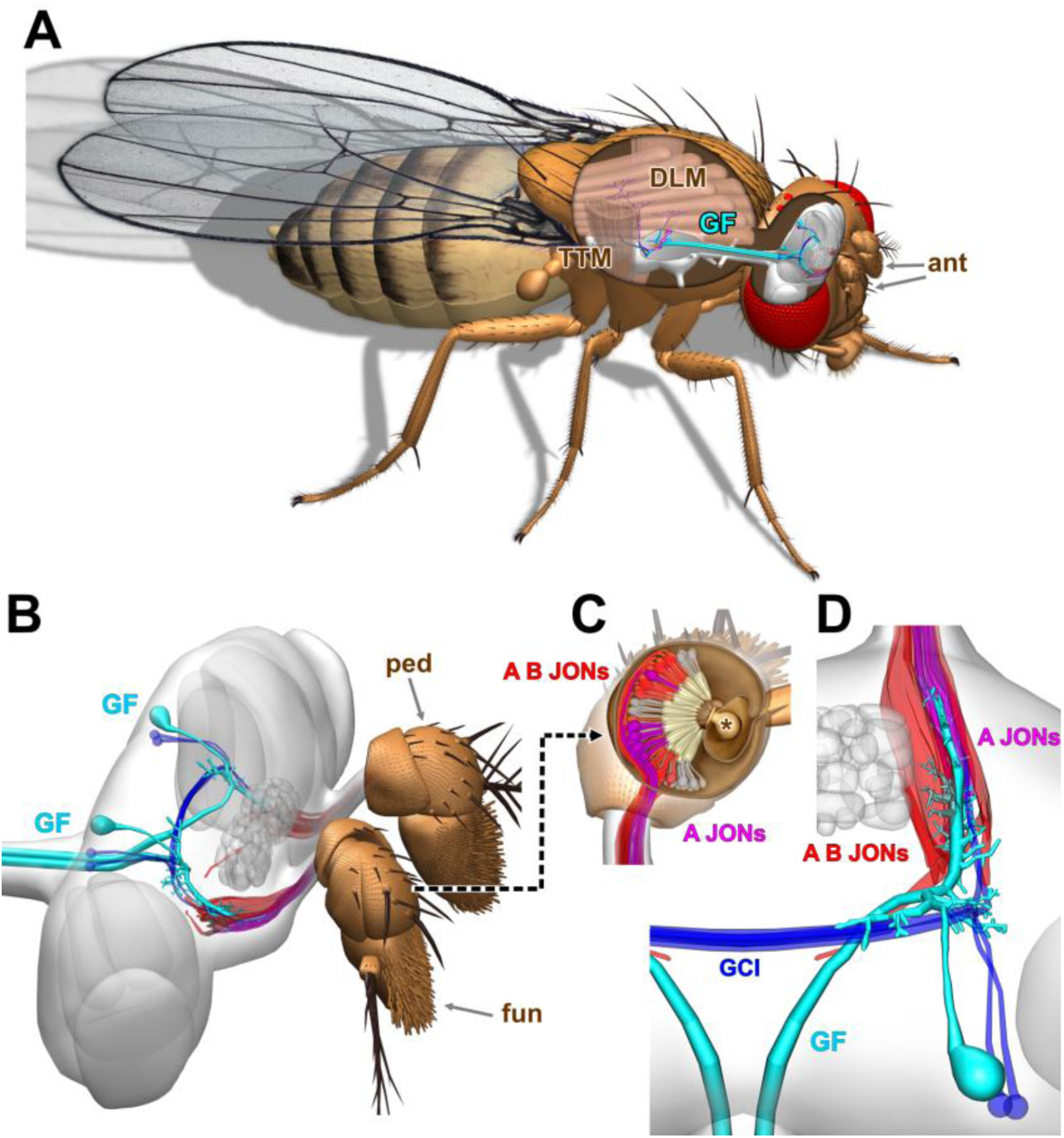
Johnston’s Organ inputs onto the *Drosophila* giant fiber escape circuit. (A) Overview of the giant fiber (GF) escape circuit, showing the GF axons descending from the brain to the thoracic ganglion, where they contact neurons that excite the tergotrochanteral ‘jump’ muscle (TTM) and the dorsal longitudinal indirect flight muscle (DLM). The antennae, one of the sources of input to the GF, are indicated (ant). (B) Side view of the pair of Giant Fiber (GF) neurons in the brain (cyan), showing their axons projecting posteriorly in the cervical connective to the thorax (left), and their main dendrites projecting anteriorly (right) to contact axons of Johnston’s Organ neurons (JONs: red and magenta) in the second antennal segment, or pedicel (ped). The third antennal segment, the funiculus (fun) is also indicated. (C) Dorsal view of the right pedicel with the top removed to show the JONs within, with their dendrites attached to the funicular stalk (asterisk). Two subpopulations of JONs are shown, the A and B JONs (red) and the A JONs (magenta). (D) Dorsal view of the right GF neuron, also showing the neighboring GCI neurons (dark blue). The main GF dendrite projects anteriorly within a cylindrical group of A JON axons, to which it is dye-coupled (magenta).

The GF responds to a synchronous combination of visual stimuli (ie. rapid looming) [7,10], and air movements [11] detected by the neurons of the antennal Johnston’s Organ (JO), the *Drosophila* analog of the mammalian inner ear [12]. There is strong evidence that ShakB is also required for these excitatory inputs onto the dendrites of the GF: the *shakB*^*2*^ mutation disrupts visual circuitry [13], and abolishes synaptic currents from auditory neurons in the JO [14]. We recently showed that, despite the presence of chemical contacts, transmission of the response to sound at the JO neuron (JON) – to – GF synapses is indeed primarily electrical [15] and that ShakB is also the most likely component of these electrical synapses onto the main dendrite of the GF [16].

The idea that gap junctions are involved in determining the development of synaptic connectivity is not new; ironically ShakB itself was first postulated to be a synaptic recognition molecule [17]. There is experimental evidence for this phenomenon in several systems. For example, the innexins Ogre and ShakB(N) are required for the proper formation of the histaminergic synapses that connect *Drosophila* retinal and lamina neurons [13]. In the leech, gap junction expression between identified neurons is a prerequisite for normal chemical synapse formation [18]. In mammals, both synapse elimination by motoneurons, and the maturation of olfactory synapses, are disrupted in connexin mutants [19,20].

Our previous work suggested that gap junction proteins can also affect synaptic connectivity at the JON-to-GF synapse. In the innexin study we observed that overexpression of the ‘correct’ ShakB isoform, ShakB(N+16), in a subset of JONs induces ectopic synaptic coupling, whereas expression of the ‘wrong’ isoform, ShakB(N), abolishes it entirely [16]. In an earlier paper we had shown that ectopic synaptic coupling could also be induced by overexpression of the transcription factor Engrailed (En) in those same sensory neurons – in that case we showed that it was accompanied by the *de novo* formation of chemical synapses, along with an increase in postsynaptic dendritic branching [15]. Taken together, these results suggested the possibility that both En and ShakB(N+16) are similarly able to alter the specificity of the JON-GF synaptic connection, by causing the formation of new mixed electrical and chemical synapses between inappropriate synaptic partners. Here we test this idea experimentally by assaying dye coupling, the distribution of putative active zones on the GF dendrites, and the distribution of GF dendrites themselves, comparing overexpression of the two different ShakB isoforms to that of En. We find that, contrary to our original hypothesis, only En is able to alter the distribution of chemical synapses (and GF dendrites), while ShakB overexpression affects gap junctional coupling alone.

## Materials and Methods

### Flies

*Drosophila melanogaster* flies of the following genotypes were obtained from the Bloomington Stock Center: *UAS-mCD8::GFP* (5130 or 5137), *JO15-GAL4* (6753). Other lines used were *UAS-shakB(N)* and *UAS-shakB(N+16)* {Pauline Phelan [6]}, *UAS-en* {Miki Fujioka [21]}, *UAS-brp-short-strawberry* {Stephan Sigrist [22]}. A GFP-tagged chromosome 2 balancer containing *CyO* (denoted *CyO-GFP* below) was obtained from Bruno Marie [23]. The following fly lines were constructed in the laboratory:

*UAS-brp-short-strawberry/ CyO-GFP; UAS-en*/*TM6B, Tb*^*1*^,

*UAS-brp-short-strawberry/ CyO-GFP; UAS-shakB(N+16)*/*TM6B, Tb*^*1*^,

*UAS-brp-short-strawberry/ CyO-GFP; UAS-shakB(N)*/*TM6B,*

*Tb*^*1*^, *UAS-mCD8::GFP/ CyO-GFP;JO15-GAL4/TM6B, Tb*^*1*^,

*GAL4* lines were crossed with the respective *UAS* lines and the F1 used for experiments. Flies were reared on cornmeal media and raised at 25°C and 60% relative humidity. In some cases, to increase *GAL4* activity, flies were transferred to 30°C or, to decrease it, to 20°C [24]. Adults from 3-10 days old were used for experiments.

### Dye coupling

Dissection and dye injection were performed as previously described [15]. Briefly, animals were dissected so as to expose the cervical connective and reveal the GF axons. One of the GF axons was impaled with a sharp glass microelectrode and injected with a mixture of 3% Neurobiotin Tracer (NB) (Vector Labs, SP-1120) and 3% Dextran Alexa Fluor 488 (DA488) (10,000 MW, Bioanalytical Instruments, D22910) diluted in distilled water. Injection electrodes were backfilled with 0.5 µl of dye mixture followed by 150 mM potassium chloride and had resistances of 45-60 MΩ resistance. The dye mixture was iontophoretically injected for up to 20 minutes using a continuous train of alternating 1 second square pulses of positive and negative current of ± 1-2 nA generated by a Master 8 stimulus generator (A.M.P.I., Israel) and delivered through an AxoClamp 2B amplifier (Molecular Devices LLC, CA, USA).

### Immunohistochemistry

The nervous systems from dye-injected flies were fixed in 4% paraformaldehyde in phosphate-buffered saline (PBS) for 45 minutes at 4°C, rinsed in several changes of PBS and further dissected to remove brain and ventral nerve cord. To process other flies, nervous systems were dissected in PBS, fixed in 4% paraformaldehyde in PBS for 30 minutes at room temperature and rinsed in several changes of PBS. After removal, nervous systems were processed for antibody labeling, and cleared and mounted as previously described [15]. nc82 (anti-Bruchpilot) antibody was obtained from the Developmental Studies Hybridoma Bank (DSHB) and used at a dilution of 1/20. Rabbit polyclonal anti-ShakB [6] was used at a dilution of 1/500. Goat anti-mouse secondary antibody labeled with Alexa-555, and goat anti-rabbit secondary antibody labeled with Pacific Blue (Thermofisher Scientific P-10994), were applied at 1/500 dilution. Streptavidin-conjugated Pacific Blue (Molecular Probes S11222) was used at 1/2000. Preparations were examined using either a Zeiss Pascal or Nikon Eclipse T1 A1r laser scanning confocal microscope and images were acquired at 8 bit resolution.

### Image processing and analysis

Confocal image stacks were imported into Fiji image analysis software [25,26], where they were 3D-rotated to a standard position and adjusted for optimal contrast. The large artefactual Brp-sh-Strawberry agglomerations noted in [15] were automatically removed from the red channel by applying a minimum filter (3 pixels) followed by a maximum filter (3 pixels), subtracting the result from the original, then readjusting the contrast. This reliably removed the agglomerations, which do not stain with Brp antibody, without affecting the actual Brp-positive active zones (S1 Fig).

Image quantification was similar to that described previously [15]. Using Fiji, the channels were Autothresholded using the Intermodes method for Brp, the Default method for the DA488 and NB signals, and the RenyiEntropy method for the ShakB signal. In all cases, the Ignore white, Stack, and Stack Histogram options were selected. The ROI was restricted to the JON afferent axons within the brain and the primary GF dendrite by first trimming the image in the 3D viewer. The thresholded images were binarized and the Measure Stack macro was used to quantify the total area of signal per slice. Areas of overlap of the Brp signal with the GF dendrites (putative active zones) were obtained using the AND operation in the Image Calculator (Fig. 2D). The unbiased Brp filtering method described above eliminated the spuriously large amount of overlap of Brp AND GF that was otherwise encountered at the posterior region of some of the GF arbors (S1 Fig). The GF dendritic surface area was estimated by measuring the total volume of a single-pixel outline (obtained by subtracting a minimum-1pixel-filtered image from the thresholded original) (Fig. 2E), then dividing by the pixel size (0.22 µm). Medially-projecting dendrites were selected by masking out all dendrites that fell within a boundary line that was 5 µm (23 × 0.22 µm pixels) medial from the center point of the GF dendrite (Fig. 2E). For the ShakB versus Brp staining, the GF dendrite was not visible so GFP-labeled JON axons were divided into A and B subpopulations by visual tracing and manual masking, followed by subtraction from the original image. The degree of retrograde Neurobiotin (NB) coupling from GF to JONs was measured by the mean cross-sectional area of NB signal averaged from 10 x1 µm slices centered on the anteriormost tip of the GF dendrite.

**Fig 2.**
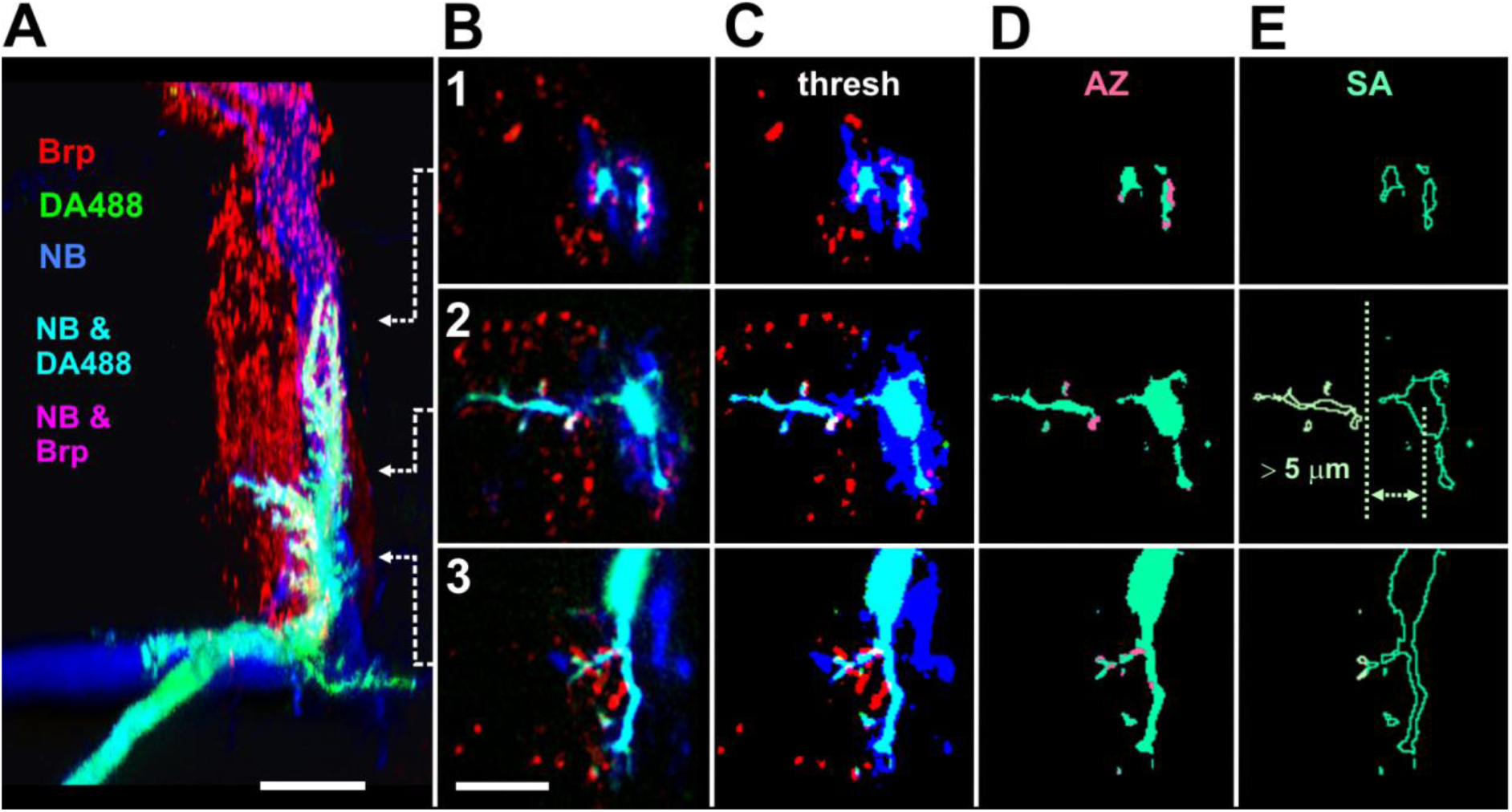
Quantification of GF synaptic inputs and branching. The GF axon was injected with a mixture of Neurobiotin (NB: blue) and Alexa Fluor 488 –coupled dextran (DA488: green), making it cyan in color. *JO15-Gal4* was used to drive expression of Strawberry-tagged Brp-short (Brp: red) in A and B subgroups of JON axons, which labels putative active zones at chemical synapses. Blue NB passes anterogradely into the A subgroup of JON axons, so their active zones appear magenta, or white when apposed to the GF. (A) Dorsal view of the right GF dendrite and Brp labeling in the A and B JON axons, in the same orientation as Fig. 1D. The subsequent panels represent single frontal slices taken from the antero-posterior levels indicated by the arrows: 1, at the anterior end of the GF dendrite, 2, midway along the dendrite in the region of medial branches, and 3, at the posterior end of the dendrite, where it bends dorsally and extends a ventral branch. (B) Single confocal slices taken at the indicated regions. Regions of overlap of Brp signal with the cyan GF dendrites (ie. putative synaptic contacts) appear white. (C) Thresholded signal (D) NB signal removed to show only putative active zones (AZ: pink) on the GF. The total AZ volume over the length of the GF dendrite was subsequently quantified. (E) Single pixel outline of the GF dendrites, used as an approximation for surface area. Dendrites projecting medially more than 5 um were masked (pale green) and quantified separately. Scale bars: 20 µm in A, 10 µm in B-E.

Maximum intensity z-series projections created in ImageJ were imported into Adobe Photoshop for construction of figure plates. The Fluorender program was used for the 3D views [27]. Blender was used for the construction of the diagrams in Fig. 1. The GF was traced in Neutube [28] and saved as swc file. This was converted to vtk format using a Python script (swc2vtk: Daisuke Miyamoto) and then to x3d format in ParaView (paraview.org). The x3d file was then imported into Blender (blender.org) for final edits and rendering. The other 3D structures of the fly body and CNS were modeled in Blender, rendered, then layered in Photoshop. Final graphics for all figures were composed and labeled using CorelDraw.

### Statistics

N represents the number of animals. The normality of the distribution of the data sets was first determined then subsequent tests were carried out using PAST3 software [29]. To identify significant differences between means of control vs. experimental groups, normally-distributed data were compared with ANOVA followed by *post-hoc* Tukey tests. In figures, * denotes p ≤ 0.05, ** p ≤ 0.01, *** p ≤ 0.001. Plots were made with Excel and transferred to CorelDraw for construction of the graphs.

## Results and Discussion

We have shown previously that driving overexpression of the transcription factor En in the JO-A and –B subpopulations of JON afferents (using the *JO15-GAL4* line [30]) causes the formation of new chemical synaptic connections between JO-B afferents and GF, in addition to expanding the population of dye-coupled afferents, and also increasing the length of medial GF dendrites [15]. Later observations suggested that, despite its apparent dissimilarity in structure and function, overexpression of the N+16 isoform of ShakB also had an identical effect on dye coupling [16]. Our hypothesis, therefore, is that overexpression of ShakB(N+16) will also cause the formation of new (mixed) synaptic connections between JON afferents and GF, resulting in the appearance of more putative active zones in contact with the GF dendrites in addition to the NB-coupling of more axons.

### ShakB(N+16) and En increase NB coupling while ShakB(N) abolishes it

We assessed changes in dye coupling by injecting the GF axon with a mixture of the dyes Dextran Alexa Fluor 488 (DA488) and Neurobiotin (NB). We used the former instead of the previously used Lucifer Yellow [15,16] because LY passes retrogradely thorough the new gap junctions induced by ShakB(N+16), making it difficult to distinguish the GF dendrites from coupled axons [16]. The *JO15-GAL4* line was used to drive *UAS-brp-sh-strawberry* in the JO-A and JO-B populations of mechanosensory afferents, as before, in order to label putative presynaptic active zones (AZ) [22]. The cross-sectional area of NB-coupled axons was averaged from 10 × 1µm slices taken at the anterior tip of the GF dendrite, where the axons leave the antennal nerve and enter the neuropil of the antennal mechanosensory and motor center (AMMC) (Fig. 3A5-8).

**Fig 3.**
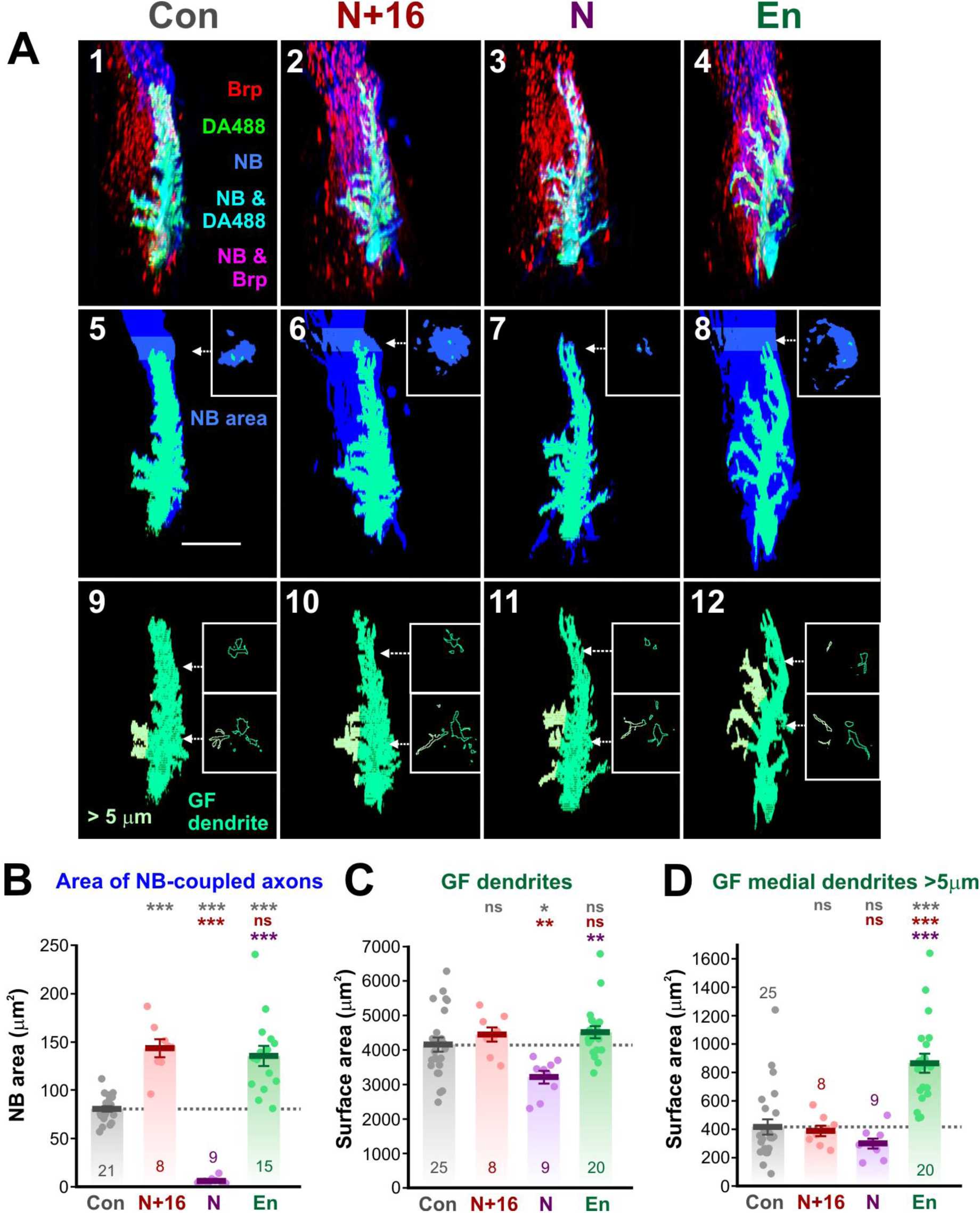
En and ShakB affect NB coupling but only En increases GF dendrites. (A) Dorsal view of the right GF dendrite from control animals (Con, column 1), *UAS-shakB(N+16)* animals (N+16, column 2), *UAS-shakB(N)* (N, column 3), and *UAS-en* (En, column 4). Panels 1-4 are typical preparations for each genotype. Panels 5-8 show the GF and NB only and indicate the 10 µm-thick region at the tip of the dendrite from which the area of NB coupling was averaged. Panels 9-12 show the surface area of the GF dendrite, with the medial dendrites projecting more than 5 µm indicated in pale yellow-green. (B-D) Scatter plots combined with bar charts showing mean ± SEM. Asterisks or “ns” above each column indicate the significance when compared to preceding columns with *post-hoc* Tukey tests. (B) Area of NB-coupled axons. Expression of both N+16 and En cause a significant increase in coupling, whereas the ShakB(N) isoform abolishes it. (C) Surface area of the GF dendrites. There are no significant increases, although *UAS-shakB(N)* expression causes a decrease. (D) Only *UAS-en* expression causes a significant increase in medial GF branching.

There is a clear 77% increase in the cross-sectional area of NB coupling with Shakb(N+16) overexpression (Fig. 3A6 and 3B). This is the same magnitude as the 68% increase in NB coupling area seen in *UAS-en* animals (Fig. 3A8, and 3B), and clearly results from the *de novo* dye coupling of a new population of JON axons, presumably the same JO-B axons, in both cases. Conversely, expression of the ShakB(N) isoform completely abolishes the normal dye coupling of the JO-A axons (Fig. 3A7 and 3B). Thus, expression of ShakB(N+16), the functional isoform at the both sides of the JO-A-to-GF synapse [16], induces the formation of NB-coupling gap junctions between JO-B axons and GF, while the non-functional Shakb(N) isoform removes or inhibits the existing ones. Whether the electrical coupling is affected remains to be tested in a future electrophysiological study.

### ShakB has fewer effects on medial GF dendrites than En

Unlike En, ShakB has less obvious effects on the surface area of GF dendrites (Fig. 3C, D). Neither ShakB(N+16) nor En alter total dendritic area, although ShakB(N) decreases it by 23% (Fig. 3C). In contrast, *UAS-en* does cause a large (108%) increase in the surface area of medial GF dendrites (defined as those that project more than 5 µm medially from the center of the GF dendrite) (Fig. 3D). This is in accordance with its effects on the measured lengths of the same dendrites, as shown previously [15]. The reduction in dendrite area in ShakB(N) animals may be due to the loss of functional gap junctions with the JO-A afferents.

### Neither ShakB(N+16) nor ShakB(N) causes an increase in chemical synapses onto GF

We have shown that ShakB(N+16) and ShakB(N) affect NB coupling with JON axons in opposite ways. The main test of our hypothesis is to determine whether the amount of presynaptic active zones (AZ) at chemical synapses is similarly affected. Again, overexpression of En was used as a positive control, where we know new putative chemical synapses are formed [15].

We estimated the numbers of putative AZ apposed to the GF dendrites by measuring the total volume of overlap of the Brp-sh and the DA488 signals. Neither the expression of ShakB(N+16) nor ShakB(N) has any significant effect. In contrast, En causes a 140% increase in putative AZ (Fig. 4B), a result consistent with our previous study [15]. Similarly, measuring AZ on the medial dendrites alone, which are expected to disproportionately bear the new synapses because the JO-B axons are medially located, shows that only En causes a large increase (of 327%). This is probably not solely attributable to the increase in medial branching (Fig.3D), because the density of AZ on the medial dendrites (calculated by dividing AZ volume by medial dendritic surface area) also increases by 102% (Fig. 4D).

**Fig 4.**
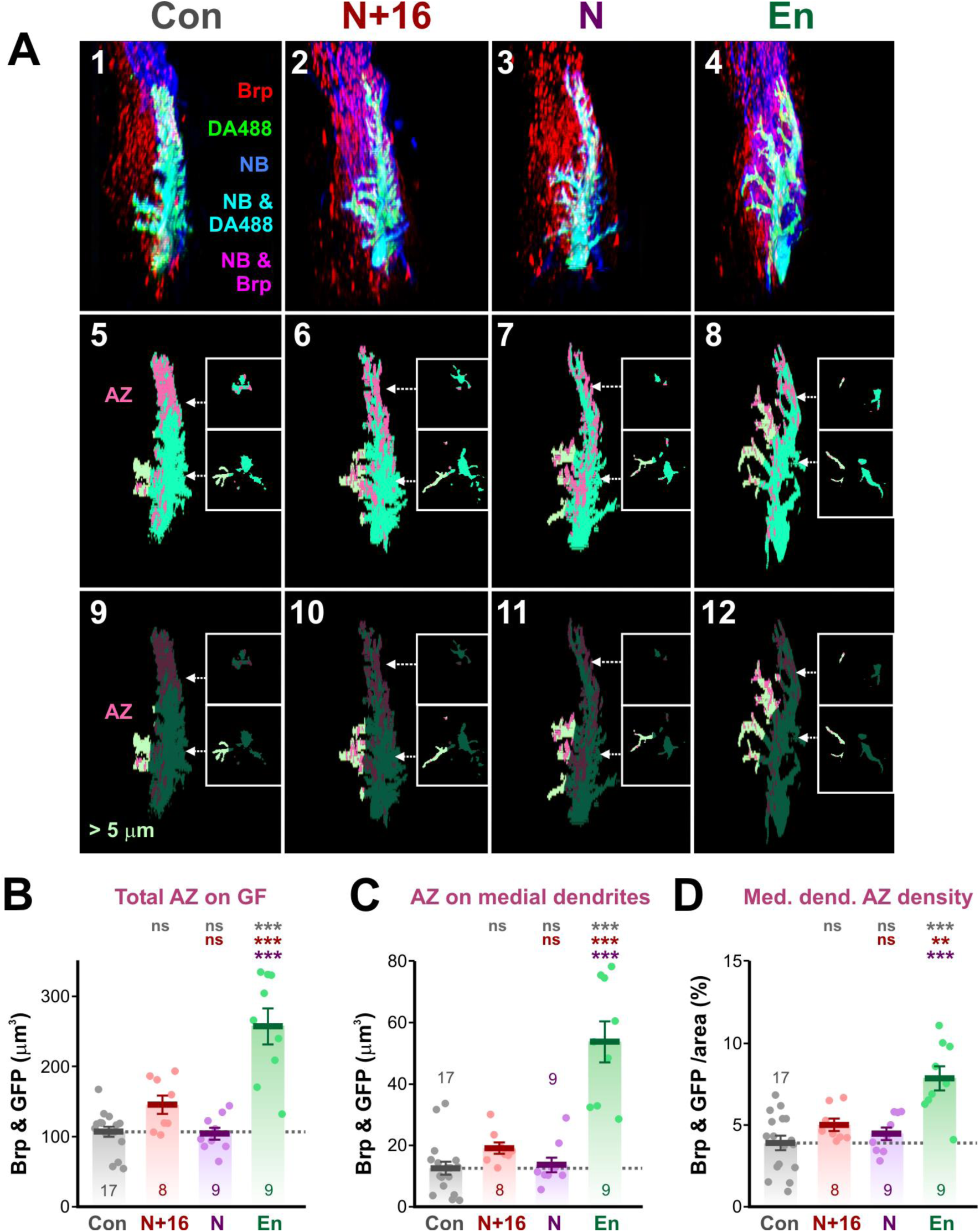
ShakB expression does not increase putative chemical synapses onto GF. (A) Dorsal view of the right GF dendrite from control animals (Con, column 1), *UAS-shakB(N+16)* animals (N+16, column 2), *UAS-shakB(N)* (N, column 3), and *UAS-en* (En, column 4). Panels 1-4 are typical preparations for each genotype. Panels 5-8 show the GF and Brp appositions (putative AZ) only. Panels 9-12 show putative AZ on only the medial GF branches. (B-D) Scatter plots combined with bar charts showing mean ± SEM. Asterisks or “ns” above each column indicate the significance when compared to preceding columns with *post-hoc* Tukey tests. (B) Total volume of AZ apposed to the GF. Only *UAS-en* expression causes a significant increase. (C) Total volume of AZ apposed to the medial GF dendrites. Only *UAS-en* expression causes a significant increase. (D) The density of AZ on the medial GF dendrites is increased by En.

This result does not therefore support our hypothesis and instead suggests that misexpression of ShakB(N+16) in JON-B axons simply results in the insertion of new gap junctions alongside chemical synapses that are already present between JO-B and GF. The physiological function of these pre-existing chemical contacts is not clear. Postsynaptic currents in GF from auditory receptor neurons are unaffected by Cd^2+^ and blocked in the *shakB*^*2*^ mutant [14], and blocking evoked transmitter release with tetanus toxin shows that they do not contribute to the GF’s response to sound [15].

### ShakB(N+16) expression increases ShakB plaques

For these experiments the GF was not dye-filled; instead we labeled the JO-A and –B axons themselves by driving *UAS-CD8::GFP* expression along with *UAS-brp-sh-strawberry* (Fig. 5). The preparations were processed for immunostaining with an antibody that recognizes all known isoforms of the ShakB protein [6]. Because the GF was not stained we did not measure the medial distance from it, instead we directly traced the JO-A and JO-B axons based on the anatomy of their arborizations [31]. ShakB immunoreactivity is typically concentrated in plaques of approximately 1 µm diameter [9,16], and these are also visible in the region of the JO-A axons that cluster around the GF dendrite (Fig. 5B).

**Fig 5.**
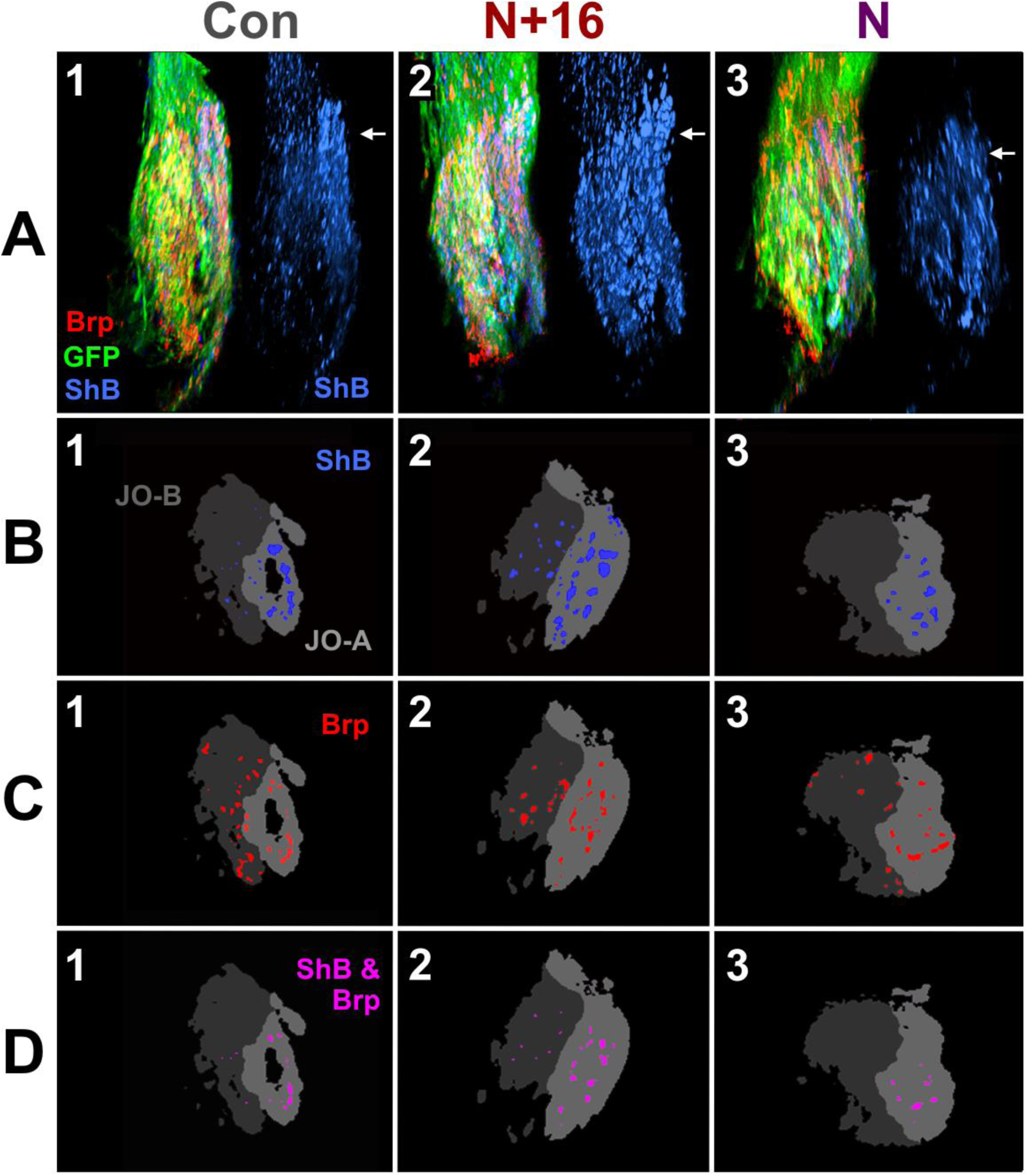
ShakB and AZ colocalization. In these preparations, *JO15-Gal4* was used to drive *UAS-brp-sh-strawberry* (Brp) and also *UAS-CD8::GFP* (GFP) in the A and B JONs. ShakB protein was stained with antibody against all isoforms (ShB). (A) Dorsal view of the right JO afferent axons from control animals (Con, column 1), *UAS-shakB(N+16)* animals (N+16, column 2), and *UAS-shakB(N)* (N, column 3). Panels 1-3 are typical preparations for each genotype. To the right of each is the ShB channel alone for clarity. Arrows indicate the approximate position of the single frontal slices shown in subsequent panels B-D. (B) Plaques of ShB labeling in the two subsets of axons, JO-A (light gray) and JO-B (dark gray). (C) Putative AZ (Brp) in the JO-A and JO-B axons. (D) Overlap of ShB and Brp in the JO-A and JO-B axons.

In controls, there is approximately 4-fold more ShakB immunostaining in the region of the JO-A axons than in the JO-B axons (Fig. 5B1 and 6A). This correlates with the normal lack of dye coupling of the GF with the JO-B axons. With expression of *UAS-shakB(N+16)* in both JO-A and –B neurons, the amount of ShakB immunostaining is greatly increased in both populations (Fig. 5B2 and 6A). This is consistent with the formation of new gap junctional plaques that allow increased dye coupling to these axons, in accordance with previous evidence that it is ShakB(N+16) that is the functional component of the JON-GF electrical synapses [16].

Conversely, no change in ShakB immunoreactivity is seen when *UAS-shakB(N)* is expressed (Fig. 5B3 and 6A). This indirectly confirms that the increase in staining seen with ShakB(N+16) expression is indeed due to the appearance of new gap junctional plaques and not simply to an non-specific accumulation of the protein in the axons – if the latter were the case we would expect an increase with ShakB(N) expression as well since the antibody recognizes both. This result also suggests that the ShakB(N) isoform, which is apparently not required for the functioning of the synapse [16], is unable to promote the assembly of additional gap junctional plaques. It does not, however, appear to reduce ShakB staining below control levels, and therefore does not completely inhibit the formation of gap junctions; those remaining presumably include the native ShakB(N+16) protein. However, ShakB(N) does have a dominant negative effect on NB-coupling, completely preventing it even though ShakB plaques remain. The sequences of ShakB(N) and (N+16) are identical except for the eponymous 16 amino-acids on the N-terminus of the latter, which is intracellular [32]. The N-terminus of ShakB is known to be important for voltage sensitivity, and also probably for trafficking and oligomerization [33]. One possibility is the presynaptic formation of heteromeric connexons composed of native ShakB(N+16) and ectopic ShakB(N), which stain with the ShakB antibody but, when docked with their homomeric ShakB(N+16) counterparts in the GF membrane [6], have an altered configuration that prevents dye coupling.

### ShakB(N+16) increases ShakB colocalization with Brp

The existence of mixed synapses at the JON-GF connection implies that the ShakB gap junction protein and the Brp AZ protein should colocalize, at least within the limits of resolution of the confocal microscope. Electron microscopy of the mixed output synapses of the GF in the thoracic ganglion showed that AZ and the distinctive large gap junctions are interspersed in close proximity, often within 0.5 µm of each other [9]. Comparable high-resolution electron microscopic data on the JON-GF connection are not available, however, examination of open-access electron microscope serial sections of the adult brain [34] show that AZ and nearby areas of close membrane apposition (a prerequisite for the existence of gap junctions) are also common between the morphologically identifiable JO-A axons and the GF dendrite (S2 Fig). However, gap junctions *per se* are not clearly visible, perhaps because the 4 nm per pixel resolution of these images is too low.

Puncta of Brp-sh fluorescence (putative AZ) are present in both the regions of JO-A and JO-B axons (Fig. 5C1). This suggests that the JO-B axons form chemical synapses but does not necessarily mean that these are with the GF itself. However, as shown above (Fig. 4), dendritic branches of GF which project too far medially to be in the region of JO-A axons do bear Brp puncta, suggesting that the JO-B axons can form chemical synapses with GF in the absence of gap junctions. In the JO-A axons however, Brp puncta tend to colocalize with ShakB (Fig. 5D1). Taken together these results suggest that the JON-GF synapse is not obligatorily mixed but can consist of either gap junctions with chemical contacts or chemical contacts alone.

As a second test of a determinative effect of ShakB on chemical synaptic connections, we assayed for co-localization of the ShakB protein with Brp puncta. Overexpression of ShakB, of either isoform, has no effect on the amount of Brp-sh labeling (putative AZ) in the JO axons (Fig. 5C2, 3 and Fig. 6B). This is an independent confirmation of the previous results of Fig. 4C that disprove our original hypothesis. Colocalization of AZ and ShakB increases with ShakB(N+16) expression (Fig. 6C), but the percentage of ShakB that is colocalized with AZ does not change (Fig. 6D). Thus the probable explanation is that colocalization becomes more frequent simply because more ShakB plaques are present, not because there is some instructive link between insertion of gap junctions and the formation of chemical synapses.

**Fig 6.**
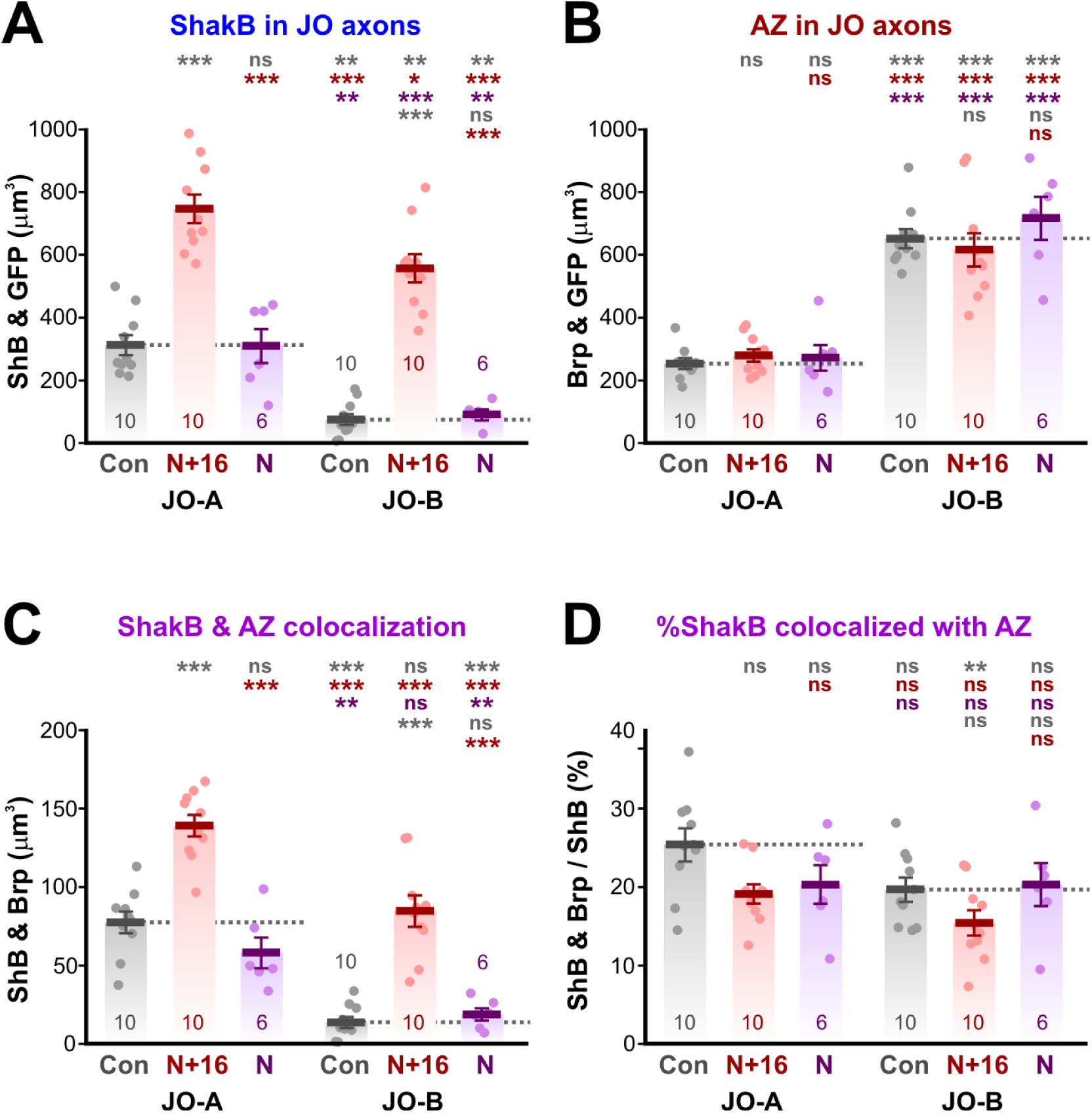
Quantification of ShakB and AZ colocalization. Quantification of data illustrated in Fig. 5. (A-D) Scatter plots combined with bar charts showing mean ± SEM. Asterisks or “ns” above each column indicate the significance when compared to preceding columns with *post-hoc* Tukey tests. (A) Total volume of ShakB staining in the two subsets of JO axons. The N+16 isoform increases ShakB staining in both JO-A and JO-B axons. (B) Total volume of putative AZ in the JO axons. More are present in JO-B than JO-A but ShakB isoform expression has no effect in either. (C) Overlap of ShakB and AZ is significantly increased by the N+16 isoform in both A and B JO axons. (D) The proportion of ShakB that is colocalized with AZ does not change.

### No evidence for instructive role of ShakB in the formation of chemical synapses

The results of this study disprove our initial hypothesis – we find no evidence that manipulating the numbers of functional gap junctions (as measured by dye coupling and by ShakB immunostaining) has any effect on the distribution of chemical synaptic connections (as measured by putative active zones). This cannot be due to the timing of expression of the *JO15-GAL4* driver because En, driven by the same driver, does increase chemical contacts. This is perhaps not an unexpected result because, even in *shakB*^*2*^ mutants, in which ShakB(N+16) and ShakB(N) are absent, the chemical portion of the GF-TTMn synapse is still present [9], and is able to partially compensate functionally [8,35]. Thus, we conclude that JO-A neurons already express ShakB(N+16) and so form gap junctions alongside their chemical counterparts, but JO-B axons only do so when made to express ShakB(N+16) artificially.

The physiological function of the chemical synapses between JO axons and GF is obscure, because it has previously been shown that blocking them with Cd^2+^ or tetanus toxin has no effect on either the postsynaptic currents in GF or its response to sound [14,15]. Morphological synapses with no discernible physiological function (ie. “silent synapses”) are in fact a common phenomenon in all nervous systems [36]. At the *Drosophila* neuromuscular junction conditionally silent Ib-type synaptic sites are important in short-term plasticity [37,38]; additionally, synaptic sites with no evoked release can evince miniature neurotransmission, which is particularly important during development [39,40]. It is a possibility that the JON-GF chemical synapses fulfil one or both of these functions. Finally, our conclusion that JON axons can form chemical presynapses with GF dendrites that have no discernible physiological function has a cautionary relevance for connectomics studies that are based on ultrastructure alone.

## Acknowledgements

The authors declare no competing financial interests. This work was supported by NINDS grant SC1NSS081726 and NIGMS grant SC3GM121190 to JMB. The Institute Pascal confocal microscope was funded by NSF DBI 0115825, DoD 52680-RT-ISP and NIMHD G12 MD007600 (RCMI). The Nikon confocal microscope was funded by NSF DBI 1337284. The Institute *Drosophila* resource center was supported by NIMHD MD007600 (RCMI). We thank Jonathan Bacon for donating the ShakB antibody. We thank Pauline Phelan and Bruno Marie for donating fly stocks. Stocks obtained from the Bloomington Drosophila Stock Center (NIH P40OD018537) were used in this study. This work was made possible in part by software funded by the NIH: Fluorender: An Imaging Tool for Visualization and Analysis of Confocal Data as Applied to Zebrafish Research, R01-GM098151-01.

## Supporting Information Captions

**S1 Fig.**
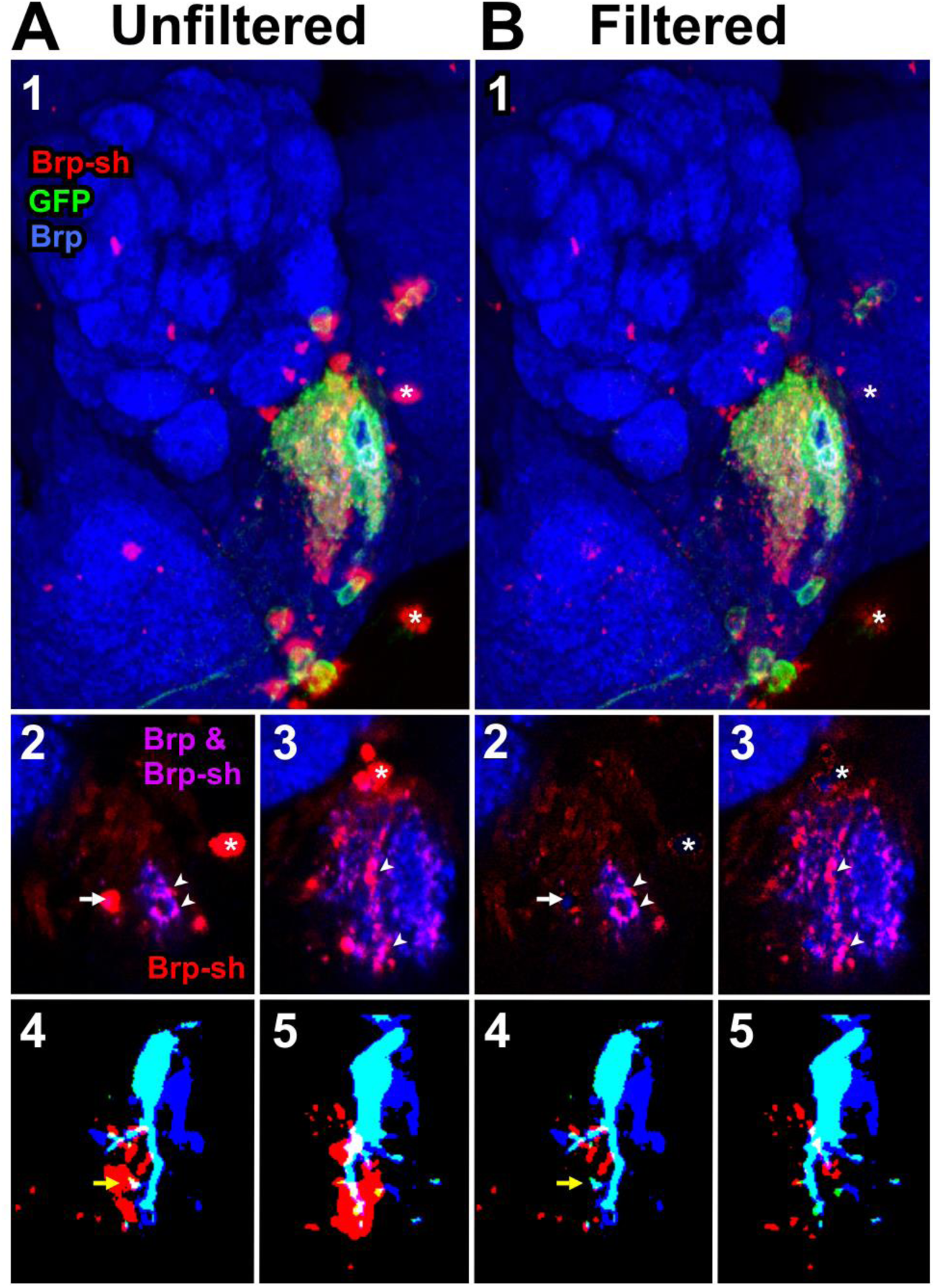
Image filtering to remove artefactual Brpsh agglomerations. The A column shows unfiltered images, the B column shows images with the red channel filtered to remove large agglomerations. Panels 1-3 show a preparation in which *JO15-GAL4* was used to drive UAS-brp-sh-strawberry (red: Brp-sh) and also UAS-CD8::GFP (green: GFP) in the A and B JONs. Native Brp protein was stained with nc82 antibody (blue: Brp). Panel1 is a maximum intensity Z-series projection through the length of the JON afferents, panels 2 and 3 are single slice views taken at the anterior tip and midway along the arbors, respectively. In these the GFP channel is removed for clarity. (A) Unfiltered images, showing large blobs that do not stain with nc82 (asterisks), some of which are associated with JO15-labeled cell bodies or axons. (B) Filtered images showing the almost complete removal of the artefactual blobs, leaving the nc82-stained Brp-sh puncta unaffected (arrowheads in 2 and 3). In panels 4 and 5 are shown preparations in which the GF axon was injected with NB (blue) and DA488 (green), making it cyan in color. Single sections are taken from the posterior region of the GF arbor. Panel 4 shows the same preparation and section as in Fig 2C3 – in this case filtering out the large blobs has little effect on the amount of overlap of Brp with the GF (white regions), although one contact region is greatly reduced (yellow arrow). In contrast, panel 5 shows the same region of a different animal in which the Brp blobs are closer to the GF dendrite so that their elimination greatly reduces the amount of spurious Brp-GF overlap.

**S2 Fig.**
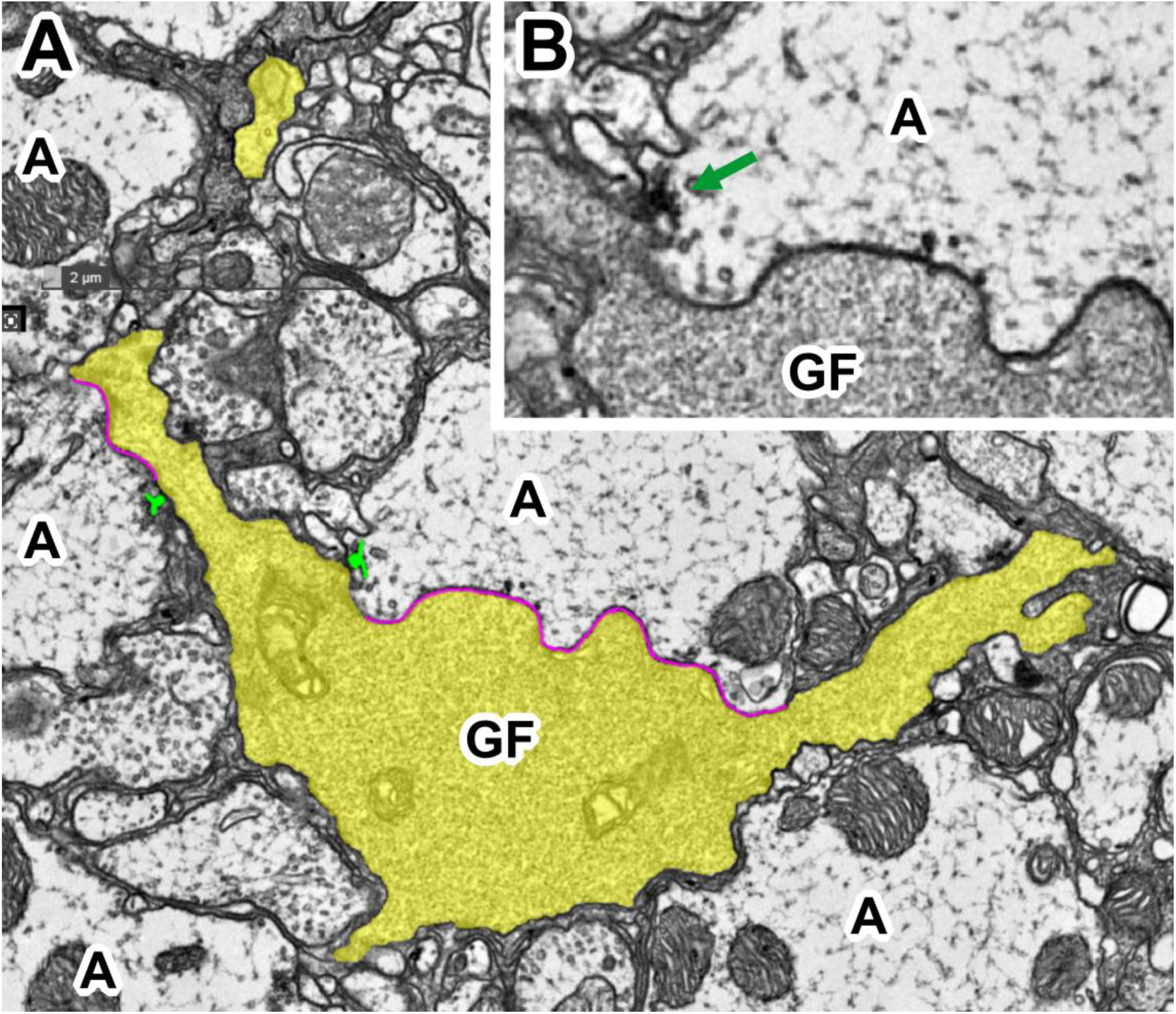
Electron microscopy of JON-GF connection. (A) Sample electron microscope section from the publicly available dataset of Tobin et al, (2017) [34]. Section number 566, location x88624, y73927, zoom level 1. The GF dendrite (GF) is colored pale yellow, areas of close apposition with JO-A axons (A) delineated with a magenta line, and chemical synapse AZ colored green. (B) The inset shows a higher magnification view without false color, and the AZ indicated with a green arrow. The resolution (4 nm per pixel) is not sufficiently high enough to unequivocally identify gap junctions in the area of close apposition, although there are similarities with the figures in [9]

